# Ribosome Profiling Reveals Translational Reprogramming via mTOR Activation in Omacetaxine Resistant Multiple Myeloma

**DOI:** 10.1101/2024.04.23.590843

**Authors:** Zachary J. Walker, Katherine F. Vaeth, Amber Baldwin, Denis J. Ohlstrom, Lauren T. Reiman, Beau M. Idler, Brett M. Stevens, Neelanjan Mukherjee, Daniel W. Sherbenou

## Abstract

Protein homeostasis is critical to the survival of multiple myeloma (MM) cells. While this is targeted with proteasome inhibitors, mRNA translation inhibition has not entered trials. Recent work illustrates broad sensitivity MM cells to translation inhibitor omacetaxine. We hypothesized that understanding how MM cells become omacetaxine resistant will lead to the development of drug combinations to prevent or delay relapse. We generated omacetaxine resistance in H929 and MM1S MM cell lines and compared them to their parental lines. Resistant lines displayed decreased sensitivity to omacetaxine, with EC50 > 100 nM, compared to parental line sensitivity of 24-54 nM. To adapt to omacetaxine, H929 and MM1S exhibited an increased percentage of multi-nucleated polyaneuploid cells that led to distinct molecular mechanisms of resistance. Interestingly, both resistant lines showed a defect in oncologic potential via extended survival in a MM xenograft model. Since omacetaxine inhibits protein synthesis, we performed both RNA-sequencing and ribosome profiling (Ribo-seq) to identify shared and unique regulatory strategies of resistance. Transcripts encoding translation factors and containing Terminal OligoPyrimidine (TOP) motifs in their 5’ UTR were translationally upregulated in both resistant cell lines. The mTOR pathway promotes the translation of TOP motif containing mRNAs. Indeed, mTOR inhibition restored partial sensitivity to omacetaxine in both resistant cell lines. Primary MM cells from patient samples were sensitive to combinations of omacetaxine and mTOR inhibitors rapamycin and Torin 1. These results provide a rational approach for omacetaxine-based combination in patients with multiple myeloma, which have historically shown better responses to multi-agent regimens.

## INTRODUCTION

Multiple myeloma (MM) is a cancer of clonal plasma cells that is characterized by the mass production of antibody protein. In the 2000s, proteasome inhibitors helped revolutionize MM treatment by targeting the proteostasis of the malignant plasma cells. Proteasome inhibitors (PIs) achieve this by blocking the disposal of protein byproducts produced during sustained antibody synthesis (1, 2). Efforts to inhibit other facets of the degradation pathways of protein homeostasis have shown preclinical potential (3, 4, 5). Currently, MM remains incurable, and patients inevitably develop resistance to proteasome inhibitors and other agents used in treatment. Thus, new therapies that retain activity in patients with relapsed disease are urgently needed. Unfortunately, other than proteasome inhibitors themselves, no other strategy targeted to intracellular proteostasis has yet had clinical success in MM.

A promising alternative approach to targeting protein homeostasis in multiple myeloma is through the blockade of protein synthesis. Previously, we and others showed that the mRNA translation inhibitor omacetaxine has potent and specific anti-myeloma activity in preclinical models, including primary samples from patients that had become proteasome inhibitor refractory (6, 7). Omacetaxine is FDA-approved for the treatment for chronic myeloid leukemia and has shown potential in other hematologic malignancies (8). This drug acts at the A-site of ribosomes, blocking the initial elongation step of translation (9). Other translation inhibitors have also shown preclinical potential in MM (10, 11). However, although multiple groups have supported pursuing mRNA translation inhibition in proteasome inhibitor-refractory MM, clinical trials have not yet been pursued. Clinical development of this new strategy is warranted, and the currently incurable status of MM will foster continued focus on this approach.

To guide clinical development, we sought to evaluate what mechanisms of resistance may emerge in MM, to allow for strategies such as drug combinations that may improve response duration. Towards this, we generated two human MM cell lines resistant to omacetaxine through continuous exposure to the drug over a prolonged period. By comparing these resistant lines to their initial, drug-sensitive counterparts, we aimed to discover the cellular alterations linked to the development of resistance. Through the utilization of gene expression analysis and ribosome profiling, we identified a mechanism of resistance shared by both cell lines involving the mTOR pathway. These findings were used to rationally develop strategies for drug combinations that may maximize the clinical application of this drug class.

## METHODS

### Cell Lines and Viability Assays

Multiple myeloma cell lines NCI-H929 and MM1S were obtained from the American Type Culture Collection (ATCC) and authenticated using short tandem repeat polymorphism profile analysis by the University of Colorado Cancer Center Cell Technologies Shared Resource Core. Cells were monitored annually for mycoplasma using the MycoAlert detection kit test (Lonza). Cells were cultured at 37 °C with 5% CO_2_ in RPMI 1640 medium containing 10% fetal bovine serum (Atlas Biologics), 100 U/mL penicillin, and 100 µg/mL streptomycin (Thermo Fisher Scientific). To determine drug efficacy, drugs were printed into 96-well culture plates using the D300e Digital Dispenser (Tecan). 60,000 cells per well were incubated in triplicate for 96 h. CellTiter-Glo Luminescent Assay (Promega) was used to determine cell viability using a Glomax Multi Detection System (Promega).

### Flow Cytometry

Multiple myeloma cell lines were washed in FACS buffer (DPBS with 2% FBS) and resuspended in 1X DAPI (Thermo Fisher). Flow data was collected on a BD FACSCelesta Multicolor Cell Analyzer (BD Biosciences) equipped with a high throughput sampler. Data was analyzed using FlowJo software (BD Biosciences). Protein translation was measured by flow cytometry by labeling nascent peptides using the puromycin analog O-propargyl-puromycin (OP-Puro) from the protein synthesis assay kit (Cayman Chemical) according to manufacturer instructions.

### Cell Line Xenografts

All in vivo animal work was approved by the Institutional Animal Care and Use Committee (IACUC) at the University of Colorado. NOD-scid gamma (NSG) mice (Jackson Labs) were injected intravenously with 2 x 10^4^ H929 or MM1S parental or omacetaxine resistant MM cells. Cells were allowed to engraft, and mice were monitored for survival. In vivo results were reported using the Kaplan-Meier survival model and hazard ratios calculated using the cox proportional hazard model.

### Confocal Microscopy

Cells for microscopy were labeled with 100 nM MitoTracker-DeepRed dye and DAPI (Thermo Fisher). Cells were washed in PBS to a final concentration of 0.5 x 10^6^ cells/mL. Slides were prepared using 150 µl of cell suspension subjected to cytospin for 3 mins at 300 rpm. Slides were air-dried and then fixed in methanol for 10 mins at -20 °C. Cells were imaged on a Zeiss LSM 780 confocal microscope equipped with a Coherent Chameleon Ultra II laser.

### RNA-seq

RNA was isolated using Trizol LS and ZYMO Directzol RNA microprep kit with on-column DNase I treatment. PolyA(+) RNA was selected from 500 ng of each input sample using the NEBNext Poly(A) mRNA magnetic isolation module. The resulting polyA RNA was used as input into the KAPA RNA Hyper prep RNAseq library prep kit (Roche). Libraries were prepared with dual-index adapters, followed by quality check of the final libraries by Qubit dsDNA HS assay and Tapestation High Sensitivity D1000 ScreenTape. The resulting libraries were pooled and submitted for 2X 150bp paired end sequencing on the Illumina NovaSeq 6000 S4 platform to a depth of 30 million paired end reads per library (Novogene Corporation Inc). Transcript expression was quantified using Salmon v1.6 (12). Differentially expressed genes were determined using the R package DESeq2 and had to have a adjusted p-value < 0.05 and a fold change > 2 (13).

### Ribosome Profiling

Ribosome profiling was performed as described previously with a few modifications (14). Cells were lysed in lysis buffer containing 1ug/mL cycloheximide. Following lysis and clarification by centrifugation, total RNA was quantified using the Qubit RNA BR assay kit. A portion of the lysate was reserved as input sample for polyA selected RNASeq preparation. The remaining lysate was used as input into the footprinting reaction with 750 U RNase I (Invitrogen Ambion). The digestion was stopped with SUPERaseIN followed by recovery of monosomes with equilibrated Microspin S-400 HR (GE) columns. Trizol LS was added to the monosome-containing elution and RNA isolated with the ZYMO Directzol microprep kit with on-column DNase I treatment. The isolated RNA was depleted of rRNA using the siTOOls riboPOOLs kit for ribosome profiling. The samples were resolved on a 15% TBE-Urea PAGE gel and ribosome protected fragments (RPFs) 28-29 nt in size were extracted and recovered. RPFs were treated with T4-PNK then used as input into the Qiagen QIAseq miRNA library prep kit with provided single index adapters. The libraries were pooled and submitted for 2X 150 bp paired end sequencing on the Illumina NovaSeq 6000 S4 platform to a depth of 60 million paired end reads per library (Novogene Corporation Inc.). Adapters were trimmed with Cutadapt (15). UMIs were trimmed and collapsed using UMI-tools (16). Transcript expression was quantified using Salmon v1.6 (12). Differentially expressed genes were determined using R package DESeq2 and deemed significant if adjusted p-value < 0.05 and fold change > 2 (13). Pipeline and all code available at https://github.com/mukherjeelab/2024_oma-resistance.

### Gene Expression Knockdowns

ON-TARGETplus siRNA SMARTPools for target genes MAPK10, FUBP1, XBP1S, and ON-TARGETplus Non-Targeting scramble control pool were purchased from Horizon Discovery. SMARTPools were resuspended at 5 mmol/L in molecular grade water. 1.5 x 10^5^ cells were combined with SMARTPools in Buffer R and electroporated using the Neon Electroporation Transfection System (Thermo Fisher) with the optimized settings of 1,200 V, 20 ms, and 3 pulses. Knockdown efficiency of siRNAs was determined by quantitative real-time reverse transcriptase PCR. RNA from electroporated cells was extracted using the RNeasy Plus Mini Kit (Qiagen) and concentration were determined with a NandoDrop system (Thermo Fisher). cDNA was generated from 50 ng RNA with the iScript cDNA Synthesis Kit (BioRad). Relative gene expression was determined using the Quanta SYBR Green Master Mix Kit (Quanta Bio) on a LightCycler 96 (Roche Diagnostics), using GAPDH as the reference gene.

### Multiple Myeloma Patient Samples

Multiple myeloma bone marrow aspirates were obtained according to the Declaration of Helsinki from patients at the University of Colorado Cancer Center following Institutional Review Board approval and written informed consent. Primary MM cells from patients were assessed for their sensitivity to drugs using the My-DST platform as previously described (17). Briefly, isolated bone marrow mononuclear cells were cultured in specific growth medium supplemented with interleukin-6. Cells were combined with drug in 96-well plate format and incubated for 48 h. Cells were washed, stained, and analyzed for viability via multi-parameter flow cytometry.

### Statistical Analysis

Unless otherwise specified, figures and statistics were generated using Prism 10 (GraphPad Software). Comparisons of two means were done using two-tailed Student *t* tests. Multiple comparison between more than two conditions was performed used ANOVA with Tukey correction. ∗p < 0.05; ∗∗p < 0.01; ∗∗∗p < 0.001; ∗∗∗∗p < 0.0001; ns, not significant. Synergy scores were calculated using SynergyFinder online tool (https://synergyfinder.fimm.fi/).

## RESULTS

### Omacetaxine Resistant MM Cells have a Unique Phenotype

Multiple myeloma cell lines H929 and MM1S were treated with escalated weekly doses of omacetaxine until they had acquired resistance compared to their isogenic parental lines, EC50s increased to > 100 nM after 96 h of treatment (Figs. 1A-B). These cell lines are referred to herein as H929-OR (Omacetaxine Resistant) and MM1S-OR. Previously, we found that baseline protein synthesis rate correlated with ex vivo omacetaxine response in primary MM samples (7). OP-puromycin staining of omacetaxine-resistant cells revealed a significant decrease in H929-OR cells, and an increase in a subpopulation of MM1S-OR (Supplemental Fig. 1A). To further identify any phenotypic differences, key MM cell surface and intracellular proteins were analyzed using mass cytometry (CyTOF). H929 parental and resistant cells were phenotypically similar via UMAP analysis, while MM1S cells formed two distinct clusters containing both parental and resistant cells (Supplemental Fig. 1B). MM1S resistant cells shifted towards a cluster containing elevated XBP1S and CD38 levels (Supplemental Fig. 1C). Together these results show the establishment of omacetaxine resistant myeloma cell lines with changes to their baseline translation rates and modified protein expression.

**Figure 1.**
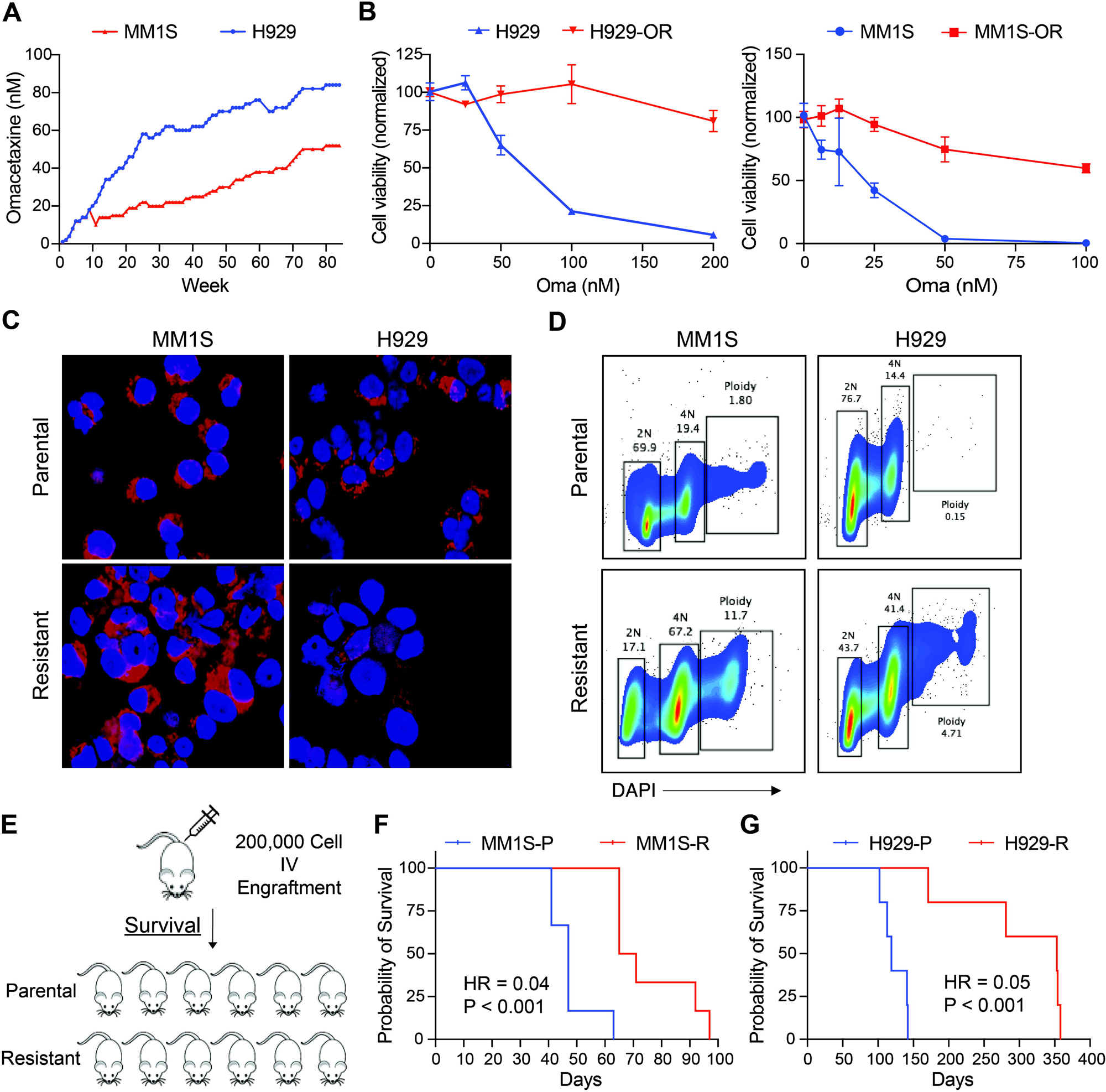
Omacetaxine Resistant MM Cell Lines have Distinct Phenotype from Isogenic Parental Cell Lines. (A) Resistant cell lines were generated by escalating omacetaxine from 1 to 84 nM in H929 and 1 to 52 nM in MM1S over the course of 70 weeks while monitoring cell viability. (B) Dose response curves of omacetaxine treated parental and resistant MM cell lines after 96 h incubation. (C) Confocal microscopy of H929 and MM1S cells stained with DAPI (blue) and MitoTracker (red). (D) Flow cytometry plots of H929 and MM1S cells stained for DAPI cell cycle. (E) Mouse model of MM parental and resistant cell lines. (F-G) Kaplan-Meier survival curves of H929 and MM1S parental vs resistant untreated xenograft model in NOD scid gamma mice (n = 5/group).

### Polyaneuploid Omacetaxine-resistant Myeloma Cells have Decreased Fitness

During routine viability checks of the resistant cells using trypan blue staining, we observed many of the cells had irregular nuclei and increased cell size compared to the parental lines. We suspected that omacetaxine exposure had resulted in polyaneuploid cancer cells (PACCs). PACCs play a role in a tumor’s ability to adapt and evolve in stressful environments including those resulting from therapeutic pressure (18). Myeloma PACCs were visible via confocal microscopy in both H929-OR and MM1S-OR by their irregular size and multi-nucleation (Fig. 1C). Increased genomic content was visible via DAPI staining and conventional flow cytometry with 8N chromatin visible in 11.7% of resistant cells compared to just 0.05% of parental in MM1S and 4.71% up from 0.15% in H929 (Fig. 1D). The increased genomic content in PACCs can result in chromosomal rearrangements, gene duplications, and numerous epigenetic changes (19).

PACCs have the ability to exit the cell cycle while under therapeutic pressure and enter brief states of reversible senescence (20). To test if the fitness of our MM cells was decreased by sustained omacetaxine exposure we used an in vivo mouse xenograft model. Two study arms were used for each H929 and MM1S comparing parental vs resistant without the presence of omacetaxine (Fig. 1E). Mouse survival was significantly longer after engraftment of the resistant line MM1S-OR compared to parental MM1S with medial survival of 68 days versus 47 days for the parental MM1S (HR = 0.04, p = 0.0006, Fig. 1F). Similarly, mice engrafted with H929-OR survived 353 versus 119 days for the parental H929 (HR = 0.05, p = 0.0018, Fig. 1G). Overall, reduced in vivo fitness of omacetaxine resistant cells may reflect that prolonged protein synthesis inhibition selects for PACCs that harbor a less robust, more indolent myeloma phenotype.

### Common and Distinct Transcriptomic Responses to Omacetaxine Resistance

To understand the gene expression changes underlying the phenotypic differences with acquired omacetaxine resistance, we next performed RNA sequencing (RNA-seq). Strong correlation within replicates (r ∼ 0.99) confirmed the high quality of the data (Fig. 2A, Supplemental Fig. 2A). There was less correlation between MM1S and MM1S-OR cells than between H929 and H929-OR cells, which suggests more omacetaxine dependent expression differences in MM1S. We found 1044 more differentially expressed genes between MM1S and MM1S-OR cells (1837) compared to H929 and H929-OR cells (793) (Fig. 2B, Supplemental Fig. 2B, Supplemental Table 1). While there were far more differentially expressed genes in MM1S cells than H929, the magnitude of expression changes was not significantly different (p = 0.08119 using a Kolmogorov Smirnov test) (Fig. 2C). Therefore, omacetaxine resistance led to a greater number of differentially expressed genes in MM1S cells, but not significantly larger changes in expression.

**Figure 2.**
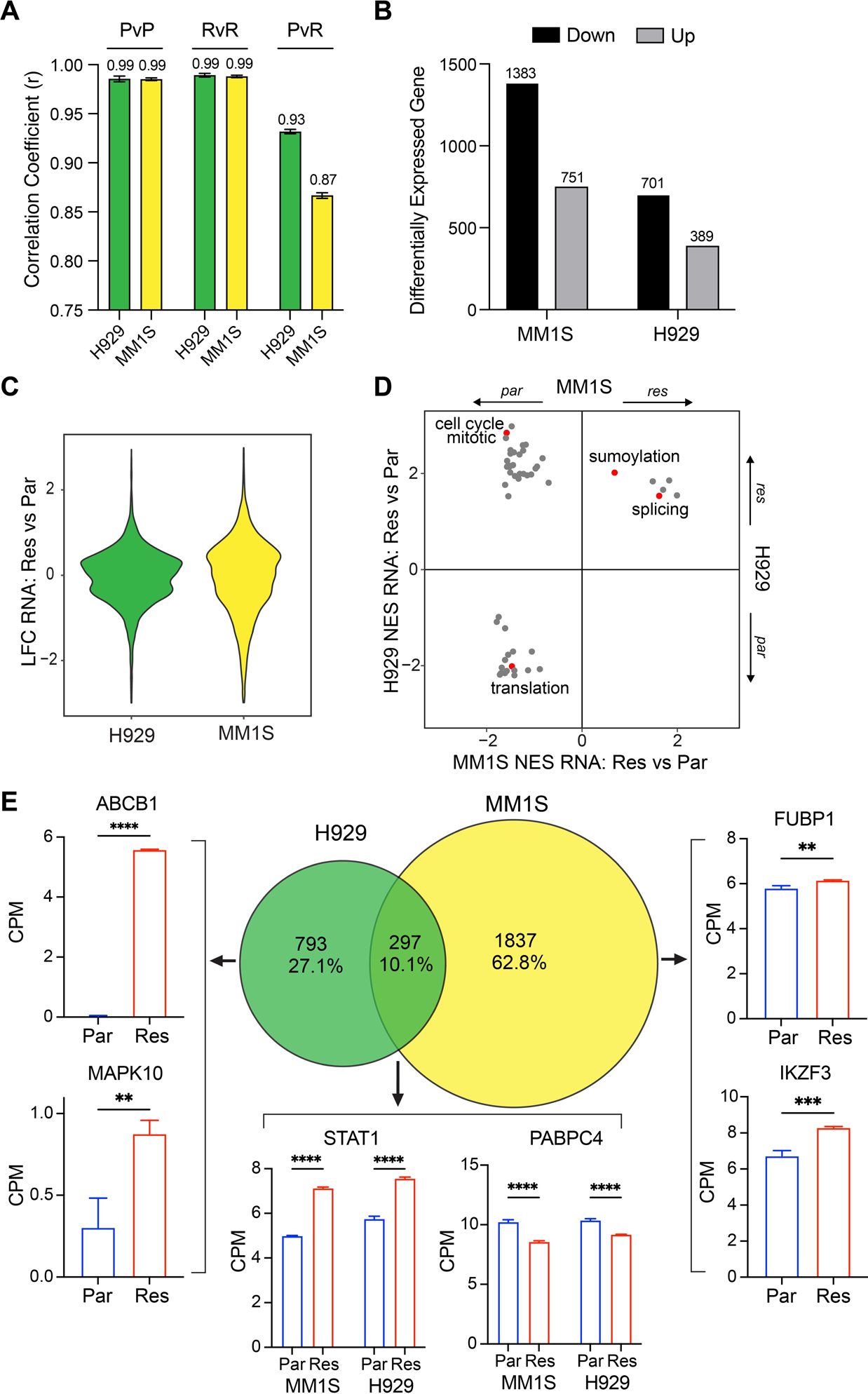
Gene Expression Changes in Omacetaxine Resistant Cell Lines. (A) Bar plot of Pearson correlation coefficients of expression within each cell line by omacetaxine resistance status, P = parental, R = resistant. (B) Bar plot of the number of statistically significant omacetaxine-dependent expression changes (adjusted p-value < 0.05 and fold-change > 2) and the direction in each cell line. (C) Violin plot of the omacetaxine-dependent log2 fold-changes in expression for each cell line. (D) Scatterplot comparing GSEA-calculated normalized enrichment score (NES) in H929 cells (y-axis) to MM1S cells (x-axis) for statistically significant gene sets (fdr < 0.05) in either cell. (E) Venn diagram of the overlap in differentially expressed genes between cell lines along with expression levels in normalized counts per million (y-axis) for individual genes of interest.

We next identified pathways that were either commonly upregulated or downregulated in both omacetaxine resistant cell lines or cell-type specific. Multiple pathways representing interferon signaling and mRNA splicing were upregulated in both MM1S-OR and H929-OR, while hemostasis and lysosome pathways were downregulated (Fig. 2D). Both omacetaxine-resistant cell lines downregulated the levels of mRNAs encoding translation factors and ribosomal proteins. We also identified a few discordantly regulated pathways related to the cell-cycle. Specifically, genes involved in prometaphase to anaphase were upregulated in H929-OR cells but downregulated in MM1S-OR cells.

Next, we identified common and unique omacetaxine induced expression changes between both cell lines. Consistent with our earlier results, we observed more than twice as many differentially expressed genes between MM1S and MM1S-OR than H929 and H929-OR (Fig. 2E). Two such genes of interest were FUBP1 and IKZF3. FUBP1 (Far upstream element binding protein 1) is a known oncogene involved in RNA splicing and MYC regulation (21, 22). IKZF3 is a myeloma promoting transcription factor involved with IMiD response (23). Two upregulated genes unique to H929-OR included ABCB1 and MAPK10. ABCB1 encodes the ATP-dependent drug efflux pump P-gp, which is commonly upregulated during the development of chemotherapy resistance in MM (24). MAPK10 is a component of the RAS/MAPK pathway involved with proliferation, differentiation, apoptosis, and transcription regulation. Expression changes shared by both cell lines included genes related to translation regulation such as PABC4. Together these results highlight shared translation pathway downregulation, as well as distinct gene expression changes caused by omacetaxine resistance in multiple myeloma cell lines.

### Differential Translation Efficiency with Omacetaxine Resistance

Given the mechanistic action of omacetaxine as a translation inhibitor, we next measured transcriptome-wide changes in translational regulation using ribosome profiling (Ribo-seq). Briefly, Ribo-seq is a method in which the mRNA footprints bound by ribosomes are purified and sequenced (25). We observed high correlation in ribosome protected fragments between replicates (average r > 0.9). Similar to RNA-seq, the lowest correlation was between MM1S and MM1S-OR cells (Fig. 3A). To determine if MM1S cells indeed had more translational changes, we calculate the translational efficiency (TE) of both MM1S and H929 cells. TE is the ratio between ribosome footprints and RNA levels and changes in this ratio represent translational regulation that cannot be identified by RNA-seq alone. MM1S cells had over 10-times more differentially translated mRNAs than H929 cells (Fig. 3B, Supplemental Table 2). Examining mRNAs with altered TE in either cell line revealed that the magnitude changes were greater in MM1S compared to H929 (p = 1.155e-14) (Fig. 3C), which was not the case for mRNA expression changes. In other words, omacetaxine resistance led to more and larger changes in translation in MM1S cells than H929 cells.

**Figure 3.**
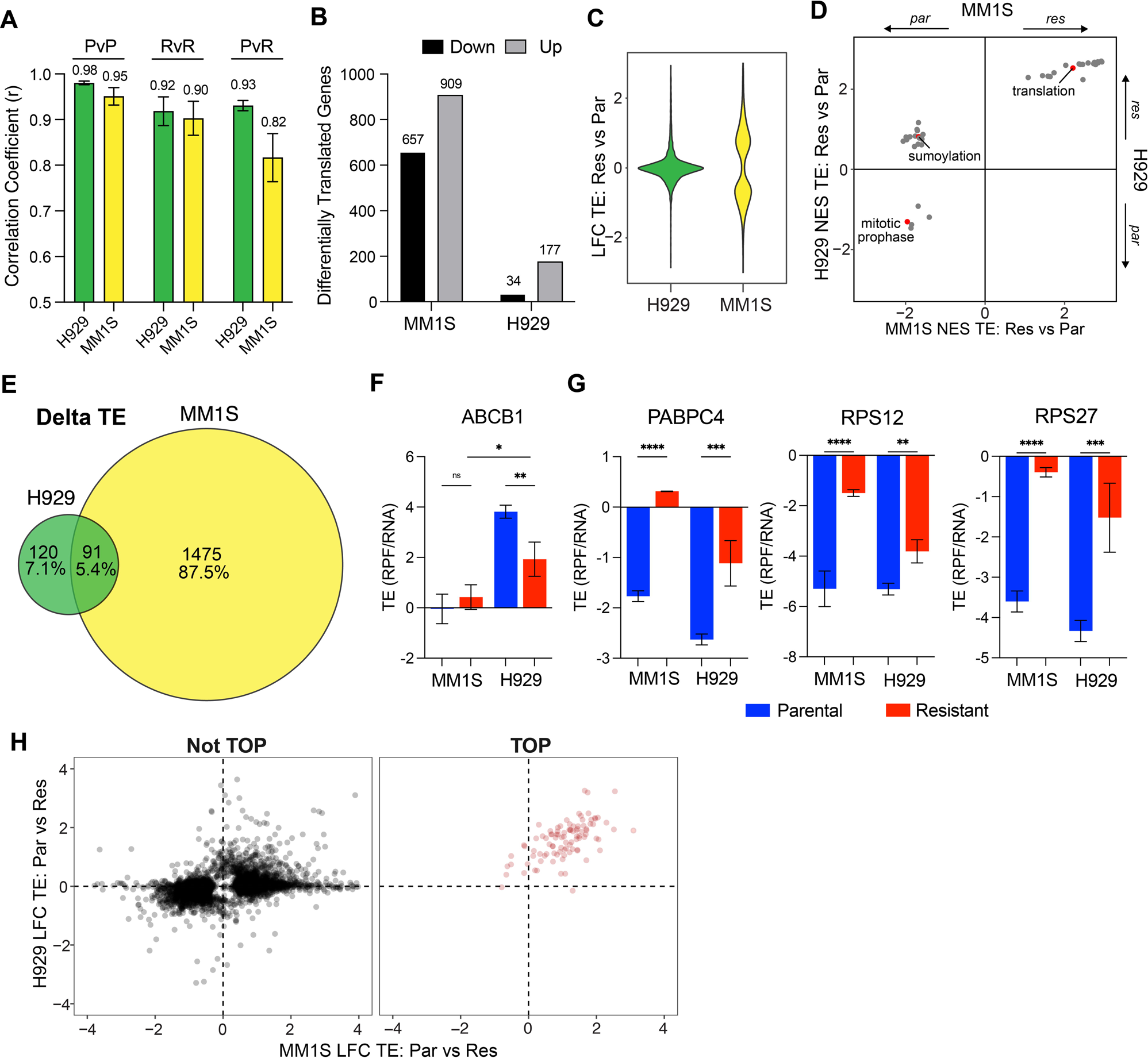
Omacetaxine Resistance Driven Translational Reprogramming in MM Cell Lines. (A) Bar plot of Pearson correlation coefficients of RPFs within each cell line by omacetaxine resistance status, P = parental, R = resistant. (B) Bar plot of the number of statistically significant omacetaxine-dependent translational changes (adjusted p-value < 0.05 and fold-change > 2) and the direction in each cell line. (C) Violin plot of the omacetaxine-dependent log2 fold-changes in translation efficiency for each cell line. (D) Scatterplot comparing GSEA-calculated normalized enrichment score (NES) in H929 cells (y-axis) to MM1S cells (x-axis) for statistically significant gene sets (fdr < 0.05) in either cell. (E) Venn diagram of the overlap in differentially translated genes between cell lines along. (F-G) Bar plot of translational efficiency changes (y-axis) for individual genes of interest. (H) Scatterplot of omacetaxine resistance driven log2-fold change in TE for mRNAs that do not contain TOP sequences (left, black) and those that contain TOP sequences (right, red) in H929 cells (y-axis) and MM1S cells (x-axis).

To further understand these changes in TE, we performed gene set enrichment analysis (GSEA) to identify pathways altered by omacetaxine resistance (Fig. 3D). Both resistant cell lines translationally repressed mRNAs encoding proteins involved in the G2 checkpoint and mitotic prophase. Transcripts encoding for translation factors and ribosomal protein pathways had upregulated TE in both omacetaxine resistant cell lines, which was the opposite of our observation of bulk mRNA levels. One possible explanation for this upregulation is that mRNAs encoding for translation factors such as ribosomal proteins contain translational regulatory elements. Collectively, these results begin to illustrate the distinct translational regulation undertaken by cells resistant to omacetaxine.

To fully understand the implications of these pathway changes, we next looked at individual transcripts. Some mRNAs exhibited similar OR-dependent changes in both mRNA levels and translation, while others resulted in discordant transcriptional and translational regulatory alterations, which would have remained hidden without ribosome profiling. Subsequently, we focused on specific mRNAs that could explain the translational changes in both cell lines. Since there were fewer differentially translated mRNAs in H929 cells (Fig. 3E), we hypothesized that H929-OR cells have experienced a lower effective concentration of omacetaxine. This would be consistent with the observation that the TE of ABCB1 was significantly higher in H929-OR (Fig. 3F). In spite of this, both OR cell lines significantly upregulated TE for the translation factors, PABPC4, RPS27, and RPS12 (Fig. 3G). We next searched for regulatory elements within these mRNAs that could elucidate the increase in TE. Markedly, the 5’ untranslated regions (UTRs) of all these mRNAs harbor an important translational regulatory element, the Terminal OligoPyrimidine (TOP) sequence. Indeed, we found that most mRNAs with TOP elements were translationally upregulated in both cell lines compared to mRNAs without TOP elements (Fig. 3H). The translation of TOP-element containing mRNAs can be modulated in an mTOR-dependent manner. Collectively these data suggest that prolonged omacetaxine exposure leads to an mTOR-dependent translational reprogramming in multiple myeloma cells through promoting the translation of mRNAs with TOP motifs that encode ribosomal proteins.

### Exploring Therapeutic Approaches to Omacetaxine Resistant Multiple Myeloma

Having shown that MM1S-OR and H929-OR cells exhibit both shared and distinct gene regulatory differences compared to parental cells, we next aimed to investigate the potential of targeting ABCB1, MAPK10 or mTOR in omacetaxine resistant MM. First, we used the direct P-gp inhibitor Zosuquidar to test if upregulation of ABCB1 promoted omacetaxine efflux activity. Cells were cultured in the presence of omacetaxine with or without 1 µM Zosuquidar for 96 h. Partial sensitivity to omacetaxine was restored in H929-OR cells, but not in MM1S-OR (Figs. 4A-B, Supplemental Fig. 2C). Next, we investigated targeting of MAPK10 upregulation in H929-OR. We treated H929 cells with MAPK10 inhibitor SP600125 alone and in combination with omacetaxine. Direct MAP10K inhibition did not restore sensitivity to omacetaxine (Supplemental Fig. 2D), but H929-OR cells appeared sensitive to single agent SP600126 at doses over 2 μM compared to little appreciable effect on parental cells (Fig. 4C). In addition, siRNA-mediated depletion of MAPK10 induced apoptosis and decreased viability of H929-OR cells (Supplemental Fig. 2E). MM1S-OR cells did not upregulate MAPK10 mRNA nor did they respond to SP600125 (Supplemental Fig. 2F). Overall, these results validate the findings of gene expression changes in H929-OR cells and found unique sensitivity to ABCB1 and MAPK10 inhibition.

**Figure 4.**
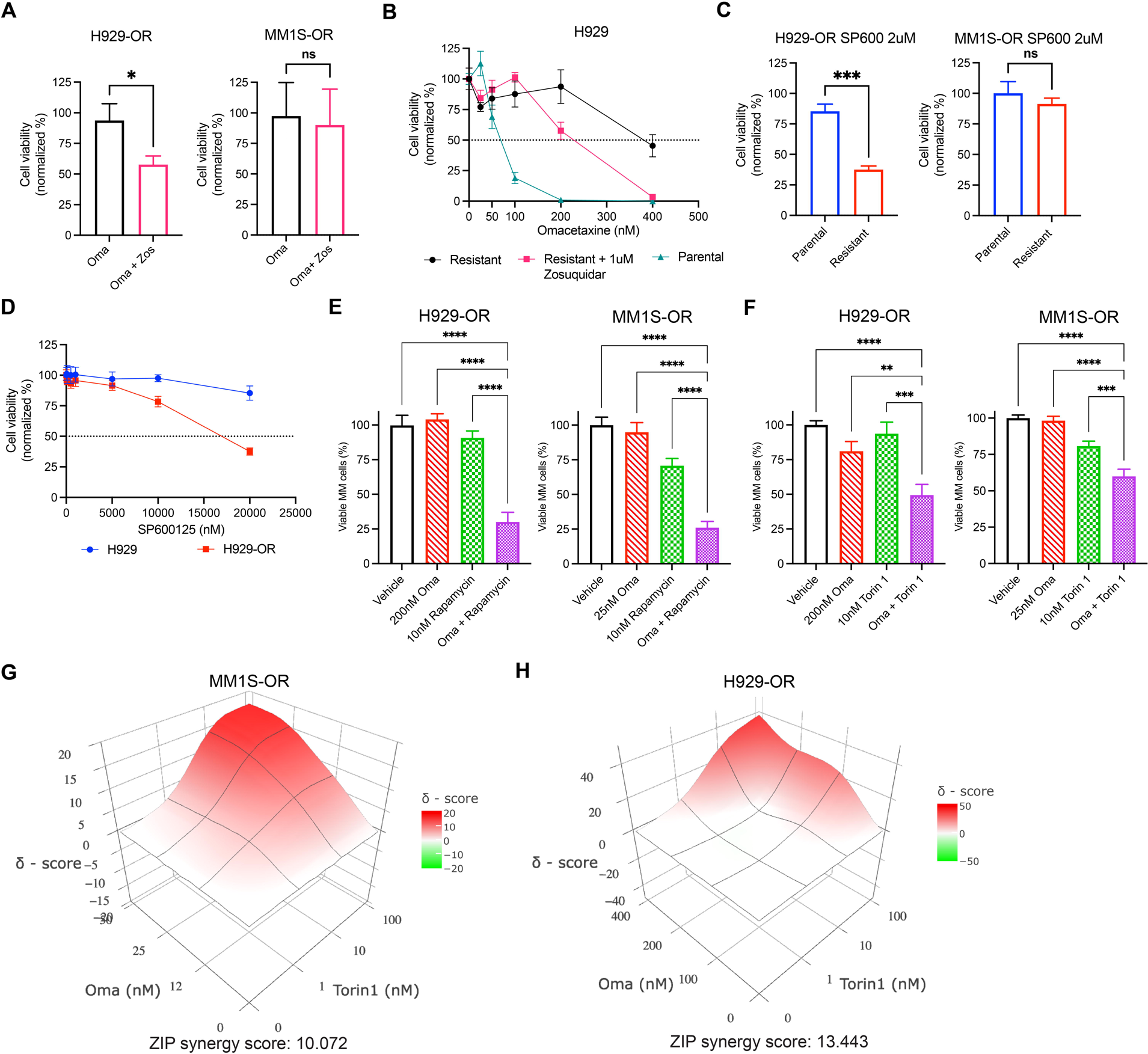
Omacetaxine Resistance Driven by Drug Efflux Pump P-gp, MAPK10, and mTOR. (A) Histogram showing 1 uM Zosuquidar restoring sensitivity to 100 nM Oma in H929-OR cells but not MM1S-OR (ANOVA p<0.05). (B) Titration of omacetaxine in H929 cells with or without the P-gp inhibitor Zosuquidar after 96 h. (C) H929-OR and MM1S-OR response to MAPK10 inhibitor SP600125 compared to parental cells after 96 h incubation. (D) Dose response curves of H929 to SP600125. (E) Response of H929-OR and MM1S-OR to single agent omacetaxine or 10 nM Rapamycin and in combination after 96 h (ANOVA p<0.001). (F) Response of H929-OR and MM1S-OR to single agent omacetaxine or 10 nM Torin 1 and in combination after 96 h. (G) Synergy plot of omacetaxine and Torin 1 in MM1S-OR cells after 96 h. (H) Synergy plot of omacetaxine and Torin 1 in H929-OR cells after 96 h. (ANOVA p<0.001). Data represent means ± SD, comparisons by two-tailed Student *t* test or two-way ANOVA with multiple comparisons (*, P < 0.05; **, P < 0.01; **, P < 0.001; ****, P < 0.0001).

MM1S-OR cells upregulated global protein translation rates and had increased TE for TOP motif containing mRNAs encoding translation factors such as ribosomal and Poly(A)-binding proteins (PABP). H929-OR cells also upregulated TE for TOP motif mRNAs. Since mTOR promotes the translation of TOP sequence containing mRNAs (26, 27, 28), we hypothesized that both omacetaxine resistant MM cells would have a common sensitivity to mTOR inhibition. To test this, the mTOR inhibitor rapamycin was titrated in the resistant cell lines. Omacetaxine resistant cells were more sensitive to 1 μM rapamycin (p = 0.0027) at 96 h (Supplemental Fig. 2G). Treatment of resistant cells with both omacetaxine and rapamycin resulted in a significant combination effect for H929-OR and MM1S-OR (Fig. 4E). Next, we tested Torin 1, a potent inhibitor of mTOR with activity against both mTORC1 and mTORC2. The 48 h combination treatment of 10 nM Torin 1 with omacetaxine in H929-OR and MM1S-OR cells had a significantly better effect than each single agent (p < 0.001) (Fig. 4F). Further combination studies revealed that omacetaxine and Torin 1 were synergistic in MM1S-OR and H929-OR with ZIP δ-scores of 10.07 and 13.44 respectively (Fig. 4G-H). Together, these results support targeting mTOR as the common dependency in omacetaxine resistant multiple myeloma cells.

### Treatment with Omacetaxine and mTOR Inhibition in Primary Multiple Myeloma

Finally, we sought to investigate if the combination of omacetaxine and mTOR inhibition was effective in killing primary multiple myeloma cells. Bulk mononuclear cells from patient bone marrow aspirates were incubated with drugs and then MM cell viability was determined by analyzing the CD38+ CD138+ population using flow cytometry (Fig. 5A). Primary sample response to single agent Torin 1 was established via dose response curve with IC50 = 2.8 μM (Fig. 5B). Single agent and combinations of low-dose omacetaxine and rapamycin resulted in significant myeloma killing in newly diagnosed patient HTB-2651 (Fig. 5C, Supplemental Table 3). We further evaluated combinations of omacetaxine and Torin 1 in both newly diagnosed and relapsed samples, yielding similar results (Fig. 5D). Overall, these data provide pre-clinical support for the rationale treatment combination of translation inhibition using omacetaxine with mTOR targeting agents in multiple myeloma.

**Figure 5.**
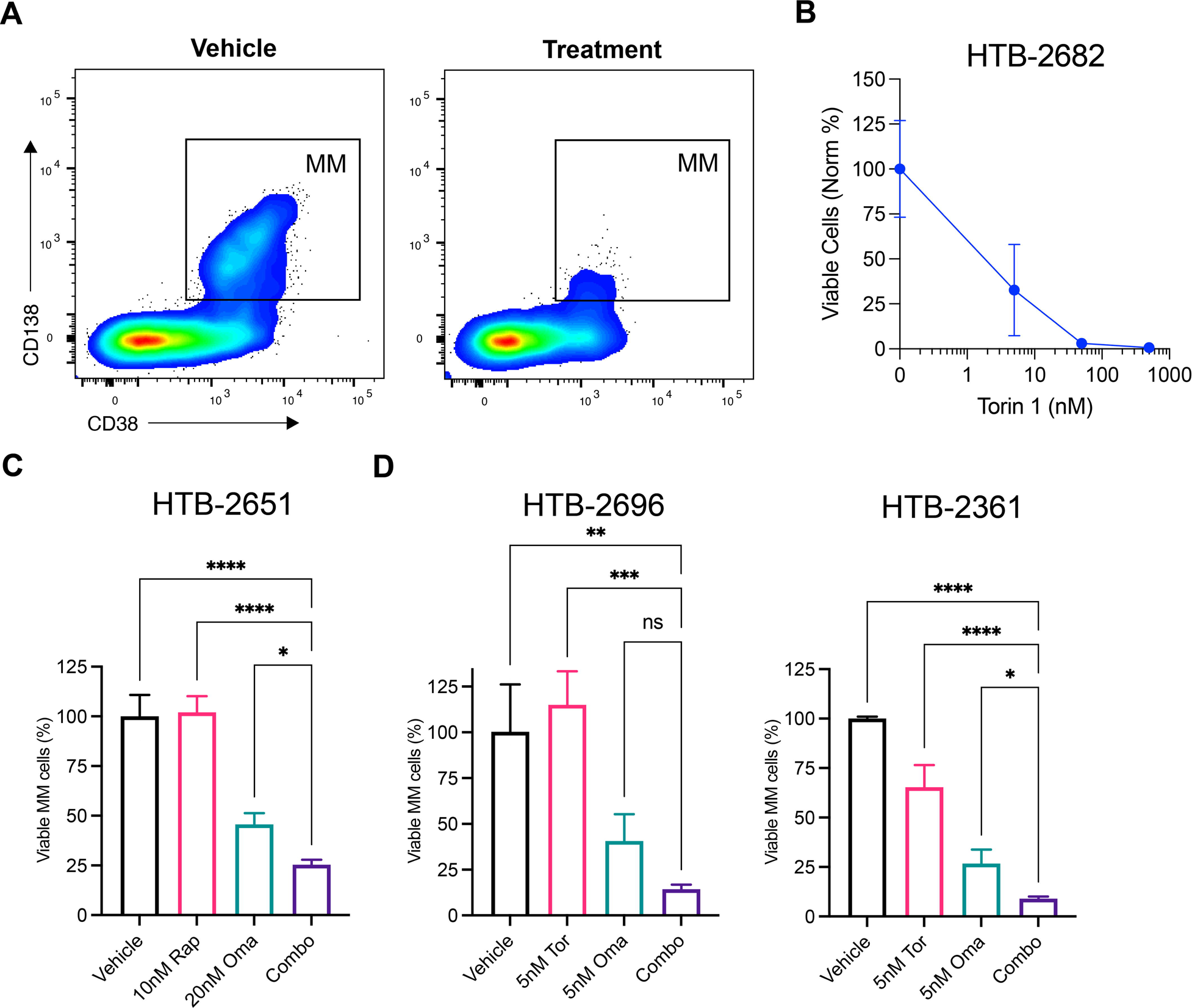
Combination Treatments with Omacetaxine and mTOR Inhibitors in Primary Multiple Myeloma Samples. (A) Flow cytometer gating strategy of primary myeloma cells after 48 h ex vivo treatment. (B) Dose-response curve of primary myeloma treated with increasing concentrations of Torin 1 for 48 h. (C) 48-hour combination treatment of rapamycin and omacetaxine in primary myeloma from newly diagnosed patient HTB-2651. (D) Combinations of Torin 1 and omacetaxine in newly diagnosed HTB-2696 and relapsed patient HTB-2361. Data represent means ± SD, comparisons by two-way ANOVA with multiple comparisons (*, P < 0.05; **, P < 0.01; **, P < 0.001; ****, P < 0.0001).

## DISCUSSION

Relapsed and refractory multiple myeloma patients continue to need new therapeutic options, even in the exciting era of T cell redirecting therapies. Patients who relapse after chimeric antigen receptor (CAR) T cells or bispecific antibodies have a dire prognosis. In this setting, treatments are often recycled from therapies used in early lines, such as proteasome inhibitors and immunomodulatory drugs. Agents with novel mechanisms of action that are distinct from those existing therapies are sorely needed. As drug combinations are more efficacious than single agents in the treatment of MM, development of new drug combinations is of particular interest, especially those that include multiple new agents that the malignant cells have not been exposed to previously. The protein synthesis inhibitor omacetaxine is a promising new backbone therapy that we have shown retains activity in multiply relapsed patients (7). The goal of this study was to better understand how MM cells respond to prolonged omacetaxine exposure and how future patients may develop resistance to this therapy. Both MM lines adapted to long-term inhibition of translation by upregulating an mTOR-dependent mechanism through the increased production of ribosomal proteins and translation factors. These resistant cells were sensitive to treatment with mTOR inhibitors rapamycin and Torin 1, a finding that translated to primary MM cells from samples dosed in combination with omacetaxine. These results provide a rational approach for combination therapy in future myeloma clinical trials involving omacetaxine.

We observed that omacetaxine resistant cells exhibited a stressed cell state. Omacetaxine resistance led to an increased unfolded-protein response, emergence of polyaneuploid cells, and a decline of in vivo fitness. The polyaneuploid response is not surprising, as this state has previously been noted after a variety of oncologics, most prominently conventional chemotherapies (19, 29). The cell line H929 harbored unique mechanisms of omacetaxine resistance compared to MM1S. The expression and translation of the mRNA encoding the classical drug efflux pump was much higher in H929 parental cells compared to MM1S parental cells and it was upregulated in H929-OR but not MM1S-OR cells. For this distinct mechanism of resistance, the difference in the initial state of ABCB1 was the likely underlying cause explaining the differential response and sensitivity to pump inhibition. In addition to ABCB1, we identified members of the MAP Kinase pathway that were highly expressed in H929-OR but not MM1S-OR cells. However, the translation of TOP sequence containing mRNAs was a common thread between both resistant cell line models, and therefore we chose to focus on this avenue. The stressed, polyaneuploid state was likely the gateway toward the selection process leading to the changes in gene expression, mRNA translational regulation, and mTOR pathway dependence that we subsequently observed in our study.

Both MM cell lines with acquired resistance to omacetaxine were differentially sensitive to mTOR inhibition via either rapamycin or Torin 1. The mTOR signaling pathway, responsible for regulating essential cellular processes like cell growth, proliferation, and survival, often exhibits dysfunction in MM (30). Due to its critical role in MM progression, the PI3K/AKT/mTOR pathway has become a promising target for therapeutic intervention. Several mTOR inhibitors have completed or are currently in clinical trials for MM treatment, however single agent mTOR trials have resulted in largely cytostatic responses (31, 32). The combination of mTOR inhibition with current therapies such as lenalidomide, dexamethasone, or bortezomib have been more promising, with up to 1/3 of trial patients receiving a partial response or better (33). Combining mTOR inhibitors with novel MM therapies such as omacetaxine represent an untouched area of clinical importance that has potential to improve outcomes for MM patients.

Our study highlights an important limitation of RNA-seq: it is only a measurement of steady-state RNA levels, but unfortunately is often misinterpreted as a surrogate for protein levels. Not only is there poor correlation between RNA and protein levels, but translational control is a dominant force in determining protein levels (34). This underlines the importance of ribosome profiling and its implications for uncovering gene regulation in cancer. Indeed, the preferential translation of TOP-motif containing RNA sequences would not have been observed in omacetaxine-resistant cells without it. This in turn led to the identification and targeting of the mTOR pathway. On the basis of this study, omacetaxine treatment combinations with mTOR inhibitors such as rapamycin or Torin 1 are rationally designed for patients with multiple myeloma. Moving forward this knowledge can be applied to the design and implementation of MM clinical trials involving either the emerging translation inhibitor drug class or to further the effectiveness of mTOR inhibitors.

## References

1. Adams J. The proteasome: a suitable antineoplastic target. Nat Rev Cancer. 2004;4(5):349–60.

2. Moreau P, Richardson PG, Cavo M, Orlowski RZ, San Miguel JF, Palumbo A, et al. Proteasome inhibitors in multiple myeloma: 10 years later. Blood. 2012;120(5):947–59.

3. Le Moigne R, Aftab BT, Djakovic S, Dhimolea E, Valle E, Murnane M, et al. The p97 Inhibitor CB-5083 Is a Unique Disrupter of Protein Homeostasis in Models of Multiple Myeloma. Mol Cancer Ther. 2017;16(11):2375–86.

4. Harnoss JM, Le Thomas A, Shemorry A, Marsters SA, Lawrence DA, Lu M, et al. Disruption of IRE1α through its kinase domain attenuates multiple myeloma. Proc Natl Acad Sci U S A. 2019;116(33):16420–9.

5. Du T, Song Y, Ray A, Wan X, Yao Y, Samur MK, et al. Ubiquitin receptor PSMD4/Rpn10 is a novel therapeutic target in multiple myeloma. Blood. 2023;141(21):2599–614.

6. Lou YJ, Qian WB, Jin J. Homoharringtonine induces apoptosis and growth arrest in human myeloma cells. Leuk Lymphoma. 2007;48(7):1400–6.

7. Walker ZJ, Idler BM, Davis LN, Stevens BM, VanWyngarden MJ, Ohlstrom D, et al. Exploiting Protein Translation Dependence in Multiple Myeloma with Omacetaxine-Based Therapy. Clin Cancer Res. 2021;27(3):819–30.

8. Novotny L, Al-Tannak NF, Hunakova L. Protein synthesis inhibitors of natural origin for CML therapy: semisynthetic homoharringtonine (Omacetaxine mepesuccinate). Neoplasma. 2016;63(4):495–503.

9. Gürel G, Blaha G, Moore PB, Steitz TA. U2504 determines the species specificity of the A-site cleft antibiotics: the structures of tiamulin, homoharringtonine, and bruceantin bound to the ribosome. J Mol Biol. 2009;389(1):146–56.

10. Li S, Fu J, Lu C, Mapara MY, Raza S, Hengst U, et al. Elevated Translation Initiation Factor eIF4E Is an Attractive Therapeutic Target in Multiple Myeloma. Mol Cancer Ther. 2016;15(4):711–9.

11. Manier S, Huynh D, Shen YJ, Zhou J, Yusufzai T, Salem KZ, et al. Inhibiting the oncogenic translation program is an effective therapeutic strategy in multiple myeloma. Sci Transl Med. 2017;9(389).

12. Patro R, Duggal G, Love MI, Irizarry RA, Kingsford C. Salmon provides fast and bias-aware quantification of transcript expression. Nat Methods. 2017;14(4):417–9.

13. Love MI, Huber W, Anders S. Moderated estimation of fold change and dispersion for RNA-seq data with DESeq2. Genome Biol. 2014;15(12):550.

14. Calviello L, Mukherjee N, Wyler E, Zauber H, Hirsekorn A, Selbach M, et al. Detecting actively translated open reading frames in ribosome profiling data. Nat Methods. 2016;13(2):165–70.

15. Martin M. Cutadapt removes adapter sequences from high-throughput sequencing reads. 2011. 2011;17(1):3.

16. Smith T, Heger A, Sudbery I. UMI-tools: modeling sequencing errors in Unique Molecular Identifiers to improve quantification accuracy. Genome Res. 2017;27(3):491–9.

17. Walker ZJ, VanWyngarden MJ, Stevens BM, Abbott D, Hammes A, Langouet-Astrie C, et al. Measurement of ex vivo resistance to proteasome inhibitors, IMiDs, and daratumumab during multiple myeloma progression. Blood Adv. 2020;4(8):1628–39.

18. Pienta KJ, Hammarlund EU, Axelrod R, Brown JS, Amend SR. Poly-aneuploid cancer cells promote evolvability, generating lethal cancer. Evol Appl. 2020;13(7):1626–34.

19. Pienta KJ, Hammarlund EU, Austin RH, Axelrod R, Brown JS, Amend SR. Cancer cells employ an evolutionarily conserved polyploidization program to resist therapy. Semin Cancer Biol. 2022;81:145–59.

20. Amend SR, Torga G, Lin KC, Kostecka LG, de Marzo A, Austin RH, et al. Polyploid giant cancer cells: Unrecognized actuators of tumorigenesis, metastasis, and resistance. Prostate. 2019;79(13):1489–97.

21. Zheng Y, Dubois W, Benham C, Batchelor E, Levens D. FUBP1 and FUBP2 enforce distinct epigenetic setpoints for MYC expression in primary single murine cells. Commun Biol. 2020;3(1):545.

22. Hoang VT, Verma D, Godavarthy PS, Llavona P, Steiner M, Gerlach K, et al. The transcriptional regulator FUBP1 influences disease outcome in murine and human myeloid leukemia. Leukemia. 2019;33(7):1700–12.

23. Bjorklund CC, Lu L, Kang J, Hagner PR, Havens CG, Amatangelo M, et al. Rate of CRL4(CRBN) substrate Ikaros and Aiolos degradation underlies differential activity of lenalidomide and pomalidomide in multiple myeloma cells by regulation of c-Myc and IRF4. Blood Cancer J. 2015;5(10):e354.

24. Besse A, Stolze SC, Rasche L, Weinhold N, Morgan GJ, Kraus M, et al. Carfilzomib resistance due to ABCB1/MDR1 overexpression is overcome by nelfinavir and lopinavir in multiple myeloma. Leukemia. 2018;32(2):391–401.

25. Ingolia NT, Ghaemmaghami S, Newman JR, Weissman JS. Genome-wide analysis in vivo of translation with nucleotide resolution using ribosome profiling. Science. 2009;324(5924):218–23.

26. Cockman E, Anderson P, Ivanov P. TOP mRNPs: Molecular Mechanisms and Principles of Regulation. Biomolecules. 2020;10(7).

27. Jia JJ, Lahr RM, Solgaard MT, Moraes BJ, Pointet R, Yang AD, et al. mTORC1 promotes TOP mRNA translation through site-specific phosphorylation of LARP1. Nucleic Acids Res. 2021;49(6):3461–89.

28. Philippe L, van den Elzen AMG, Watson MJ, Thoreen CC. Global analysis of LARP1 translation targets reveals tunable and dynamic features of 5’ TOP motifs. Proc Natl Acad Sci U S A. 2020;117(10):5319–28.

29. Gerashchenko BI, Salmina K, Eglitis J, Huna A, Grjunberga V, Erenpreisa J. Disentangling the aneuploidy and senescence paradoxes: a study of triploid breast cancers non-responsive to neoadjuvant therapy. Histochem Cell Biol. 2016;145(4):497–508.

30. Pene F, Claessens YE, Muller O, Viguié F, Mayeux P, Dreyfus F, et al. Role of the phosphatidylinositol 3-kinase/Akt and mTOR/P70S6-kinase pathways in the proliferation and apoptosis in multiple myeloma. Oncogene. 2002;21(43):6587–97.

31. Günther A, Baumann P, Burger R, Kellner C, Klapper W, Schmidmaier R, et al. Activity of everolimus (RAD001) in relapsed and/or refractory multiple myeloma: a phase I study. haematologica. 2015;100(4):541.

32. Farag SS, Zhang S, Jansak BS, Wang X, Kraut E, Chan K, et al. Phase II trial of temsirolimus in patients with relapsed or refractory multiple myeloma. Leukemia research. 2009;33(11):1475–80.

33. Ramakrishnan V, Kumar S. PI3K/AKT/mTOR pathway in multiple myeloma: from basic biology to clinical promise. Leuk Lymphoma. 2018;59(11):2524–34.

34. de Sousa Abreu R, Penalva LO, Marcotte EM, Vogel C. Global signatures of protein and mRNA expression levels. Mol Biosyst. 2009;5(12):1512–26.

