## Supplemental Figures for "Ribosome Profiling Reveals Translational Reprogramming via mTOR Activation in Omacetaxine Resistant Multiple Myeloma"

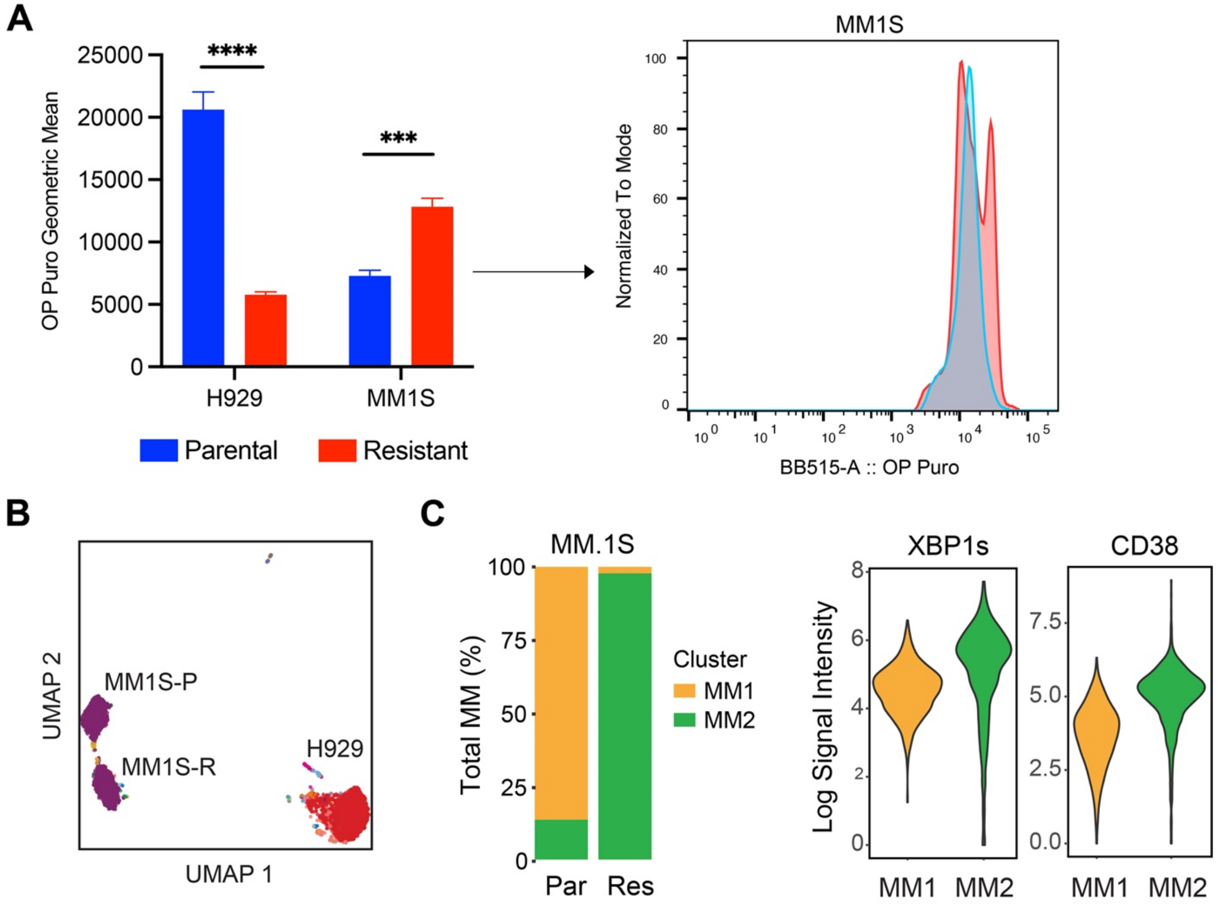


**Supplemental Figure 1. Characterization of omacetaxine resistant MM cell lines.** (A) Global protein translation rates of H929 and MM1S cells as measured using OP-Puromycin. (B) Uniform Manifold Approximation and Projection (UMAP) clustering of parental and resistant cells generated from CyTOF results. (C) MM1S clustering percent based on UMAP groups and mean expression of XBP1s and CD38 in MM1S cells determined via CyTOF.

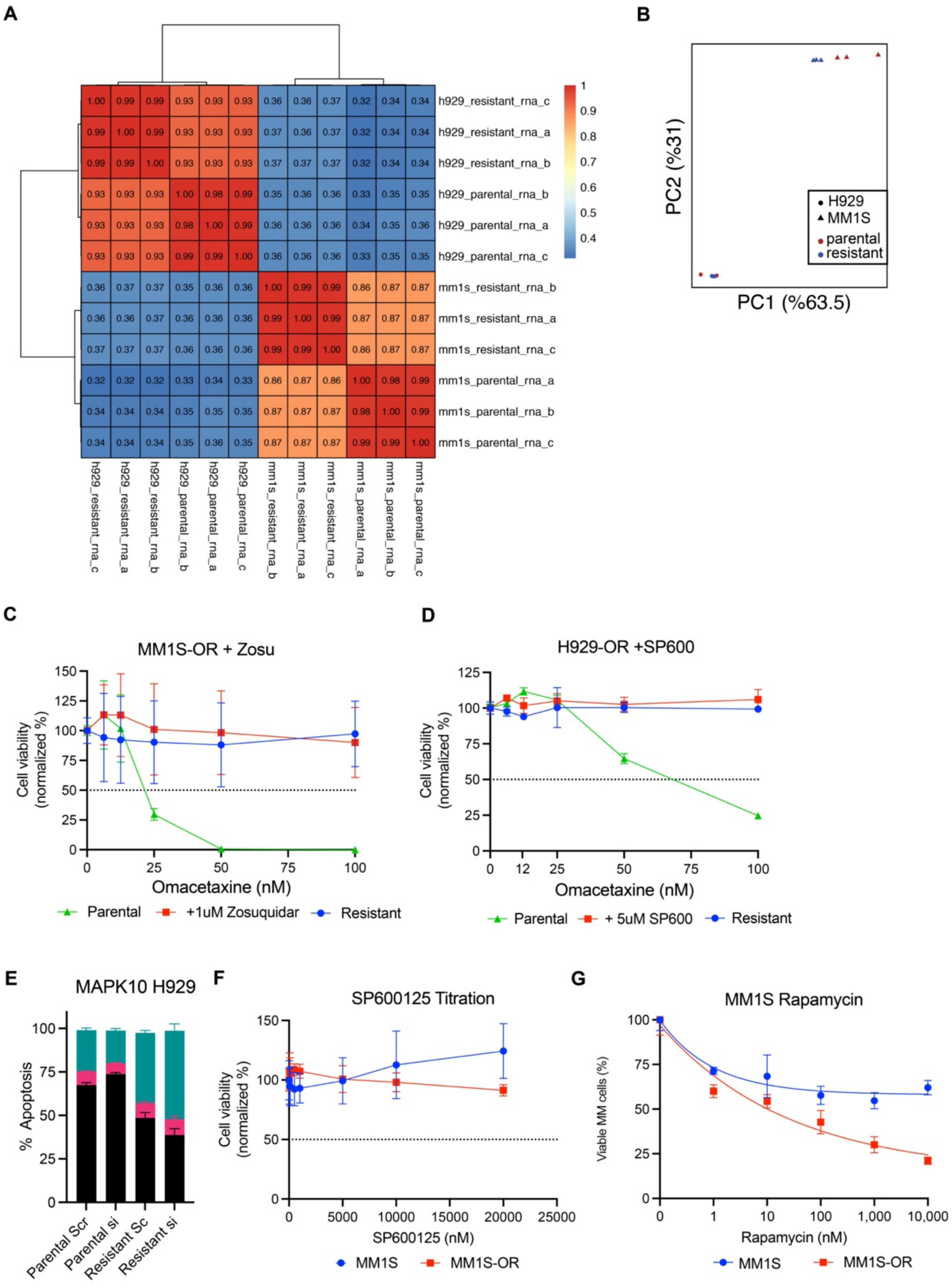


**Supplemental Figure 2. Omacetaxine resistance mechanisms in H929 and MM1S.** (A) RNAseq correlation data from H929 and MM1S cells. (B) PCA of gene expression analysis for H929 and MM1S oma resistant and parental cell lines. (C) Titration of omacetaxine in MM1S cells with or without the P-gp inhibitor Zosuquidar after 96 hr. (D) H929 cell response to combination of MAPK10 inhibitor SP600125 and omacetaxine after 96 hr incubation. (E) Apoptosis plot of H929-OR 24 hrs post siRNA electroporation with MAPK10 knockdown. (F) MM1S-OR cell response to MAPK10 inhibitor SP600125 compared to parental cells after 96 hr incubation (G) Titration of mTOR inhibitor rapamycin in MM1S cells after 96 hr incubation.

**Supplemental Table 3: Myeloma Patient Characteristics**

| ID | Age/Sex | Disease State | Cytogenetics | Treatment History |
|---|---|---|---|---|
| HTB-2682 | 35M | New Dx | Normal | N/A |
| HTB-2651 | 72F | New Dx | High Risk | N/A |
| HTB-2696 | 64M | New Dx | Normal | N/A |
| HTB-2361 | 44M | Multi-relapse | Normal | VRD, Dara, Car/Pom/Dex, Elo/Car/Pom, Car+DCEP, CAR-T |

VRD= velcade revlimid dexamethasone, Dara = daratumumab, Car = carfilzomib, Dex = dexamethasone, Pom = pomalidomide, Elo = elotuzumab, DCEP = dexamethasone, cyclophosphamide, etoposide, and cisplatin
